# Alcohol availability during withdrawal gates the impact of alcohol vapor exposure on responses to alcohol cues

**DOI:** 10.1101/2021.12.21.473655

**Authors:** M.J. Carpio, Runbo Gao, Erica Wooner, Christelle A. Cayton, Jocelyn M. Richard

## Abstract

**Background:** Chronic intermittent ethanol (CIE) vapor inhalation is a widely used model of alcohol dependence, but the impact of CIE on cue-elicited alcohol seeking is not well understood. Here, we assessed the effects of CIE on alcohol-seeking elicited by previously learned cues, and on acquisition of new cue-alcohol associations.

**Methods:** In Experiment 1, male and female Long Evans rats were first trained in a discriminative stimulus (DS) task, in which one auditory cue (the DS) predicts the availability of 15% ethanol and a control cue (the NS) predicts nothing. Rats then underwent CIE or served as controls. Subsets of each group received access to oral ethanol twice a week during acute withdrawal. After CIE, rats were presented with the DS and NS cues under extinction and retraining conditions to determine whether they would alter their responses to these cues. In Experiment 2, rats underwent CIE *prior* to training in the DS task. We also assessed alcohol consumption, aversion-resistant drinking, somatic withdrawal symptoms, and behavior in an open field.

**Results:** We found that CIE enhanced behavioral responses to previously learned alcohol cues, but only in rats that received access to alcohol during acute withdrawal. CIE disrupted cue responses in rats that did not. When CIE occurred before cue learning, male rats were slower to develop cue responses and less likely to enter the alcohol port, even though they had received alcohol during acute withdrawal. We also found that CIE increased alcohol consumption and aversion-resistant drinking in male but not female rats.

**Conclusions:** These results suggest that CIE alone does not potentiate the motivational value of alcohol cues, but that an increase in cue responses requires the potentiation of the value of alcohol during acute withdrawal. Further, under some conditions CIE may disrupt responses to previously learned and subsequently acquired alcohol cues.

## Introduction

Environmental cues that predict the availability of drugs like alcohol can drive several aspects of addiction, including escalation of alcohol use and the propensity to relapse even after long periods of abstinence (Courtney et al., 2015; Preston et al., 2018). Chronic intermittent ethanol (CIE) vapor inhalation is a commonly used model of alcohol dependence which has been shown to drive increased alcohol consumption and self-administration, including habitual and compulsive alcohol seeking behaviors (Gilpin et al., 2008; Griffin et al., 2009; Radke et al., 2017; Renteria et al., 2018; Xie et al., 2019). Yet we know relatively little about the impact of CIE vapor inhalation on reactivity to alcohol cues, including in models of relapse.

The existing literature on the impact of CIE vapor inhalation on behaviors related to alcohol cues is mixed. For instance, CIE vapor inhalation appears to enhance cue-induced reinstatement of alcohol seeking under some conditions (Liu and Weiss, 2003, 2002), but not others (Ciccocioppo et al., 2003; Eisenhardt et al., 2015). One salient difference between the methods used in these studies is the availability of alcohol during acute withdrawal: studies showing enhanced cue-induced reinstatement have included periods of access to alcohol during the vapor phase when animals are in acute withdrawal (Liu and Weiss, 2003, 2002). Importantly, one issue with the cue-induced reinstatement model of relapse is that it involves response-contingent presentations of cues. This makes the model, essentially, a test of conditioned reinforcement (i.e., the degree to which these cues can reinforce behaviors), but not necessarily the conditioned motivational properties of cues (i.e., the degree to which cue presentations can invigorate behavior). Therefore, cue-induced reinstatement may be of limited relevance to cue-driven relapse in humans, which is typically precipitated by cues outside of the individual’s control (Epstein et al., 2009; Preston et al., 2018, 2009). Additionally, reinstatement of alcohol seeking by response-contingent presentation of cues is often less robust than reinstatement induced by non-contingent contextual cues signaling alcohol availability (Tsiang and Janak, 2006). This may be why many studies investigating alcohol relapse use a combination of contingent and non-contingent cue presentations (Ciccocioppo et al., 2003, 2002; Williams and Schimmel, 2008), muddling precise interpretation of their results.

Given the issues with response-contingent cue presentation as a model of relapse to alcohol seeking, it is important to understand the effect of CIE vapor inhalation on responses to non-contingent alcohol cues. Work in this area thus far has been limited: Kufahl et al. (2011) found that CIE vapor inhalation did not alter alcohol-seeking behavior in the presence of non-contingent (olfactory and auditory) contextual cues predicting the availability of alcohol. Importantly, animals in that study did not have the opportunity to consume or self-administer alcohol in acute withdrawal during the vapor phase of this experiment. Therefore, it remains unknown whether CIE vapor inhalation can enhance behavioral responses to contextual alcohol cues, if given access to alcohol during acute withdrawal. Additionally, the effects of CIE vapor inhalation on the invigoration of alcohol-seeking behaviors by discrete cues signaling alcohol availability have not been reported.

To address these open questions, we examined the effects of CIE vapor inhalation on behavioral responses to previously learned cues signaling alcohol availability, and the effects of CIE on subsequent alcohol cue learning and responsivity. In Experiment 1 we trained rats in a discriminative stimulus (DS) task with alcohol reward (Ottenheimer et al., 2019; Richard et al., 2018) prior to CIE vapor inhalation. We then assessed the effects of CIE on alcohol consumption, somatic withdrawal signs, behavior in an open field, and alcohol cues responses under both extinction and retraining conditions. In Experiment 2 we assessed the effects of CIE on subsequent cue learning in the DS task, as well as aversion-resistant drinking. We found that the effects of CIE on alcohol cue responses depend on both the timing of CIE relative to cue learning, and on access to alcohol during acute withdrawal. Additionally, we found sex-specific effects of CIE on alcohol consumption and aversion-resistant drinking.

## Materials and Methods

### Subjects

Long-Evans rats (n= 101, 59 females and 60 males; Envigo) weighing 225-275 grams at arrival were used in this study. Rats were individually housed with *ad libitum* access to food and water and were maintained on a 14-hr/10-hr light/dark cycle (lights on at 6am; lights off at 8pm). All experimental procedures were approved by the Institutional Animal Care and Use Committee at the University of Minnesota and were carried out in accordance with the Guide for the Care and Use of Laboratory Animals (NIH).

### Discriminative Stimulus (DS) Task Training

Rats were trained in operant chambers (29.53 x 23.5 x 27.31 cm) equipped with tone and white noise generators, and a reward port with a head entry detector (Med Associates Inc., Fairfax, VT). Prior to DS task training we conducted port training sessions in which 15% ethanol (.1 mL) was delivered to the reward receptacle via syringe pump. Sessions terminated when rats entered the port after 30 successive alcohol deliveries or 120 minutes had elapsed, whichever occurred first. A second supplemental port entry training session was provided for rats that did not enter the port 30 times in the first session. The third and final port entry session included all rats. After port training, rats were trained in a DS task, as described previously (Ottenheimer et al., 2019; Richard et al., 2018, 2016). In this task, entry into the port during one auditory cue (the DS) resulted in delivery of 15% ethanol (0.1mL) to the port. Port entry in the absence of the DS or during the non-rewarded stimulus (NS) had no programmed consequences. Auditory cues were a combination of 2700 and 4100kHz tones (DS) and white noise (NS). Training consisted of five stages. In stages 1-4 30 DS cues were presented with decreasing lengths (60s, 30s, 20s, 10s). Rats advanced to the next stage when they had entered the port on at least 60% of the DS cues. The NS cue (10s) was introduced in the fifth stage and was also presented 30 times for a total of 60 cues. For all stages, cues were presented on a pseudorandom variable interval schedule with varying mean intertrial intervals to keep the sessions at 90 minutes.

### Chronic Intermittent Ethanol (CIE) Vapor Exposure

Rats underwent chronic intermittent exposure (CIE) to ethanol vapor in a passive vapor inhalation system (La Jolla Alcohol Research, Inc., San Diego, CA) that is described in detail elsewhere (Gilpin et al., 2008). During vapor exposure, rats were individually housed in standard static cages with transparent walls and *ad libitum* access to food and water. Vapor chamber settings were adjusted according to daily intoxication ratings and weekly blood ethanol concentrations (BECs). CIE was administered for 14-hours per session, 4 sequential nights a week, for 3 weeks. After exposure sessions intoxication levels were rated according to behavioral markers for both CIE and control rats, as described elsewhere (Barker et al., 2017). BECs were assessed once a week after the first vapor exposure cycle. Blood was collected from the tail vein using a 70μL heparinized capillary tube. Plasma was separated from the red blood cells, and injected into an alcohol analyser (Model AM1, Analox Instruments Ltd., Stourbridge, United Kingdom). Somatic withdrawal signs were assessed during the final week of vapor exposure, 6-8 hours after CIE rats were removed from the vapor chambers (Majchrowicz, 1975).

### Cue Test: DS Extinction and Retraining

Approximately 2.5 weeks after the completion of CIE (Experiment 1) or the end of DS task training (Experiment 2), rats underwent four cue tests in the same behavioral chambers used for DS task training. During the first cue test rats were presented with 10 probe cues (5 DS and 5 NS) under extinction conditions (no ethanol reward). Cue testing under extinction was performed to differentiate behavioral responding to the cue from behavioral changes driven directly by ethanol reinforcement. Immediately afterwards, 60 cues (30 DS and 30 NS) were presented under retraining conditions, where port entry during the DS once again resulted in an alcohol reward. Rats underwent three additional testing sessions, spaced 48 hrs apart, with increasing number of trials or amounts of ethanol delivery in the following order: 1) 80 cues (40 DS and 40 NS), 2) 120 cues (60 DS and 60 NS), and 3) 120 cues with double the amount of ethanol per reward (0.2ml), to identify maximal responding for ethanol.

### Quinine-Adulterated Alcohol Consumption Tests

To assess the degree to which the rats’ consumption of alcohol was compulsive rats underwent quinine-adulterated alcohol consumption tests before and after vapor exposure or task training (Hopf and Lesscher, 2014; Seif et al., 2013). Rats were given home cage access to 15% ethanol with 0, 10, 30 and 90 mg/L quinine hydrochloride (Seif et al., 2015; Sneddon et al., 2018), for 24-hr periods, with at least 24 hrs in between tests.

### Open Field Tests

Exploratory behavior in an open field was assessed before and after CIE during 10-minute sessions, in an open-top, white, opaque box measuring 60 x 60 x 40 cm. Behavior was videorecorded and analyzed offline. The first open field test occurred prior to CIE, the second occurred 6-8 hrs after the last vapor exposure session, and the third occurred 2 weeks after the last vapor exposure day. Video recordings were analyzed for rat position and activity using EthoWatcher software (Crispim Junior et al., 2012), which were analyzed using custom MATLAB scripts. Thigmotaxis (the tendency to remain close to walls) was measured as an index of anxiety-like behavior. Distance traveled was used as a measure of total locomotor activity.

### Experiment 1: Impact of CIE vapor inhalation on responses to previously learned alcohol cues

Rats (n=80; 40 males, 40 females) received 8 weeks of intermittent access (Hopf et al., 2010) to 15% ethanol in their home cages to acclimate them to its taste and pharmacological effects, and to ensure they were sufficiently motivated to learn the DS task for alcohol reward. Rats that failed to drink at least 2 g/kg/day during the final 2 weeks of pre-exposure were excluded from further testing (n=20; 13 males; 7 females). Rats that met drinking criteria (n=60; 33 females, 27 males) then underwent DS task training. They were trained for 15 days with the DS only, followed by 12 sessions that included the NS for a total of 27 training sessions. At this point, rats that failed to meet learning criteria (DS response probability > 0.5, NS response probability < 0.5, DS/NS response ratio > 1.5) were designated as “non-learners” and excluded from further assessment of cue-related behavior (n=17; 7 males; 10 females) but were assessed for the effects of CIE vapor inhalation on alcohol consumption and open field behavior. Rats were then assigned to either the CIE or control groups, matched and counterbalanced based on DS task performance. Prior to CIE, rats underwent assessment of initial behavior in an open field, and quinine-adulterated alcohol consumption, followed by 3 weeks of CIE as described above.

Rats that were over-exposed to ethanol vapor during this phase were excluded from further assessment (1 female, 2 males). To assess the impact of access to ethanol during acute withdrawal, some rats (n=28; 7 male controls, 6 male CIE, 8 female controls, 8 female CIE) received access to 15% ethanol in the home cage for 2-hr periods, 2 days a week, 6-8 hours after CIE rats were removed from the vapor chambers, whereas the remaining rats did not receive access to oral alcohol during the vapor exposure phase (n=30; 5 male controls, 8 male CIE, 7 female control, 10 female CIE). Following the CIE phase, we assessed behavior in an open field and quinine-adulterated alcohol consumption, as described above. Finally, rats underwent final cue testing as described above.

### Experiment 2: Impact of CIE vapor inhalation on alcohol cue learning and responsivity

Rats (n=39; 20 males, 19 females) were given 2 weeks of intermittent access to 15% ethanol in their home cages and were then assessed for quinine-adulterated alcohol consumption and behavior in an open field as described above. They were then assigned to matched CIE and control groups based on their baseline alcohol consumption (n=9-10 per group). Rats underwent 3 weeks of CIE as described above; all rats received access to 15% ethanol in the home cage for 2-hr periods, 2 days a week, 6-8 hours after CIE rats were removed from the vapor chambers. Rats that were over-exposed to ethanol vapor were excluded from further assessment (3 females). Following CIE, rats were reassessed for open field behavior and quinine-adulterated alcohol drinking as described above. Then, they underwent training in the DS task: rats were trained with the DS cue alone for 10 sessions, followed by 12 sessions with both the DS and the NS. Following an additional assessment of quinine-adulterated alcohol drinking they underwent final cue testing as described above.

### Statistical Analysis

Statistical analyses were conducted using MATLAB (Mathworks). The effects of CIE vapor exposure, sex, and access to ethanol during acute withdrawal were assessed using linear mixed models, with random effects included for subject. Fixed effects for session, test number, and/or quinine concentration were included as appropriate for each behavioral test.

## Results

### Female rats consumed more ethanol during intermittent access and DS task training

In our first experiment rats received 8 weeks of homecage intermittent access to 15% ethanol prior to training the DS task. We found that female rats consumed significantly more g/kg ethanol throughout intermittent access (Fig 1A; main effect of sex, F(1,1895)=36.798, p < 0.001), including during the final 2 weeks (Fig 1B; t(78)=2.6386, p = 0.01). Males escalated their consumption across intermittent access, but females did not (main effect of session: F(1,1844)=62.146, p < 0.001; session X sex interaction: (F(1,1844)=20.605, p < 0.001; main effect of session, males: F(1,938)=85.757, p < 0.001; females: F(1,906)=1.4385, p = 0.23). Female rats also consumed more g/kg ethanol across training in the DS task (Fig 1C; main effect of sex: F(1,1615)=19.411, p < 0.001). Increased g/kg consumption in the task may be partially due to the limit on maximal alcohol consumption during each session (30 trials x 0.1 mL 15% ethanol) and the consistent weight differences between males and females in this study (F(1,1615)=704.23, p < 0.001). Across training rats increased their port entry response probability during the DS (Fig 1D; F(1,1611)=33.237, p < 0.001) and decreased their port entry probability during the NS (F(1,703)=29.687, p < 0.001), with no differences observed between male and female rats (F values of 0.344 to 1.56). After 27 sessions rats differed in their responses to the DS and NS in both response probability (Fig. 1E; F(1,80)=704.65, p < 0.001) and response latency (F(1,57)=13.3, p < 0.001). Rats were then assigned to matched CIE or control groups based on DS task performance.

**Figure 1.**
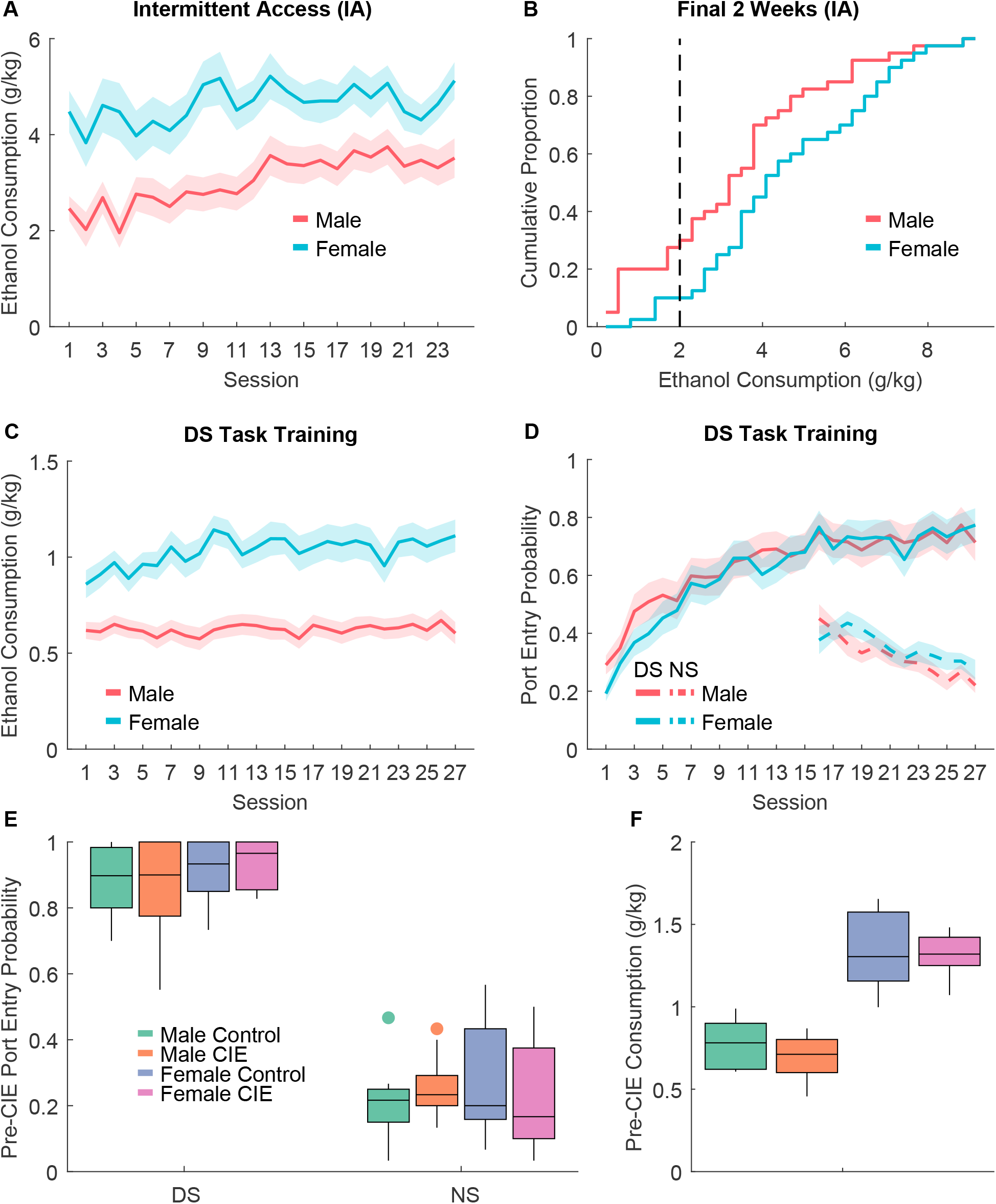
DS task training prior to vapor exposure. A) Ethanol consumption (g/kg) during homecage intermittent access, prior to task training. B) Distribution of mean ethanol consumption during the final 2 weeks of intermittent access in males (red) versus females (blue). The dotted line marks the minimum consumption required during this period to move on to training in the DS task. C) Ethanol consumption during training in the DS task. D) Port entry probability during the DS ethanol cue (solid line) and NS control cue (dotted line) across training. E) Final port entry probability during the DS (left) and NS (right) in rats assigned to vapor exposure (CIE) or control conditions after training. F) Ethanol consumption during the final DS task training session.

### CIE vapor inhalation increased behavioral responses to alcohol cues, but only in rats that received access to ethanol during acute withdrawal

Approximately 2.5 weeks after the end of CIE rats were tested for their responses to the alcohol DS and control NS cues under extinction conditions (Fig. 2A). We found that the effects of CIE on port entry probability during the DS depended on whether rats received access to alcohol during acute withdrawal (CIE X alcohol access interaction: F(1,47)=6.01, p = 0.018): CIE rats that received alcohol access were more likely to enter the port during the DS than controls on average, whereas CIE rats that did not receive alcohol access had a lower port entry probability, though no pairwise comparisons were significantly different after corrections for multiple comparisons (Fig. 2A). We found no effect of sex on DS port entry probability (main effect of sex: F(1,47)=1.12, p = 0.295; CIE X sex interaction: F(1,47)=1.36, p = 0.25; CIE X alcohol access X sex interaction: F(1,47)=0.43, p = 0.51). We also found no effects of sex, CIE or alcohol access on port entry probability during the NS (F values of 0.05 to 0.73). We next assessed whether CIE would alter response probability or latency under retraining conditions (with ethanol delivery). Because we were concerned about potential ceiling effects with the standard number of trials, we also tested rats with increasing numbers of trials or ethanol reward per trial (increasing the maximal ethanol reward per session). While female rats consumed more g/kg ethanol during these sessions (Fig. 2B; F(1,222)=36.837, p < 0.001) there was no interaction between sex and CIE vapor inhalation or alcohol access during withdrawal (F values ranging from 0.19 to 1.06), and female and male rats did not significantly differ in their response probability (Fig. 2C; main effect of sex: F(1,222)=0.2553, p = 0.61). Additionally, because adding sex as a variable did not improve the fit for our linear mixed model analysis of DS port entry probability (AIC=−199.26 without sex versus −189.7 with sex) we combined males and females for subsequent analyses. Overall, we found that increasing the trial number and reward volume reduced port entry probability (Fig. 2C; F(1,212)=4.683, p = 0.0358), but that there was no interaction between this factor and CIE or alcohol access (F values 0.155 to 1.34). When we pooled data across the tests, we found that CIE had opposing effects on DS port entry probability, depending on whether the rats had alcohol access during acute withdrawal (CIE x alcohol access interaction, F(1,216)=5.48, p = 0.020), increasing port entry probability in rats that received access, and decreasing port entry probability in rats that did not (similar to the test results under extinction conditions). Increasing the trial number and/or reward volume also decreased NS port entry probability (F(1,212)=6.96, p = 0.009). We observed no effect of CIE or interactions with alcohol access for NS port entry probability (F values 0.072 to 0.75).

**Figure 2.**
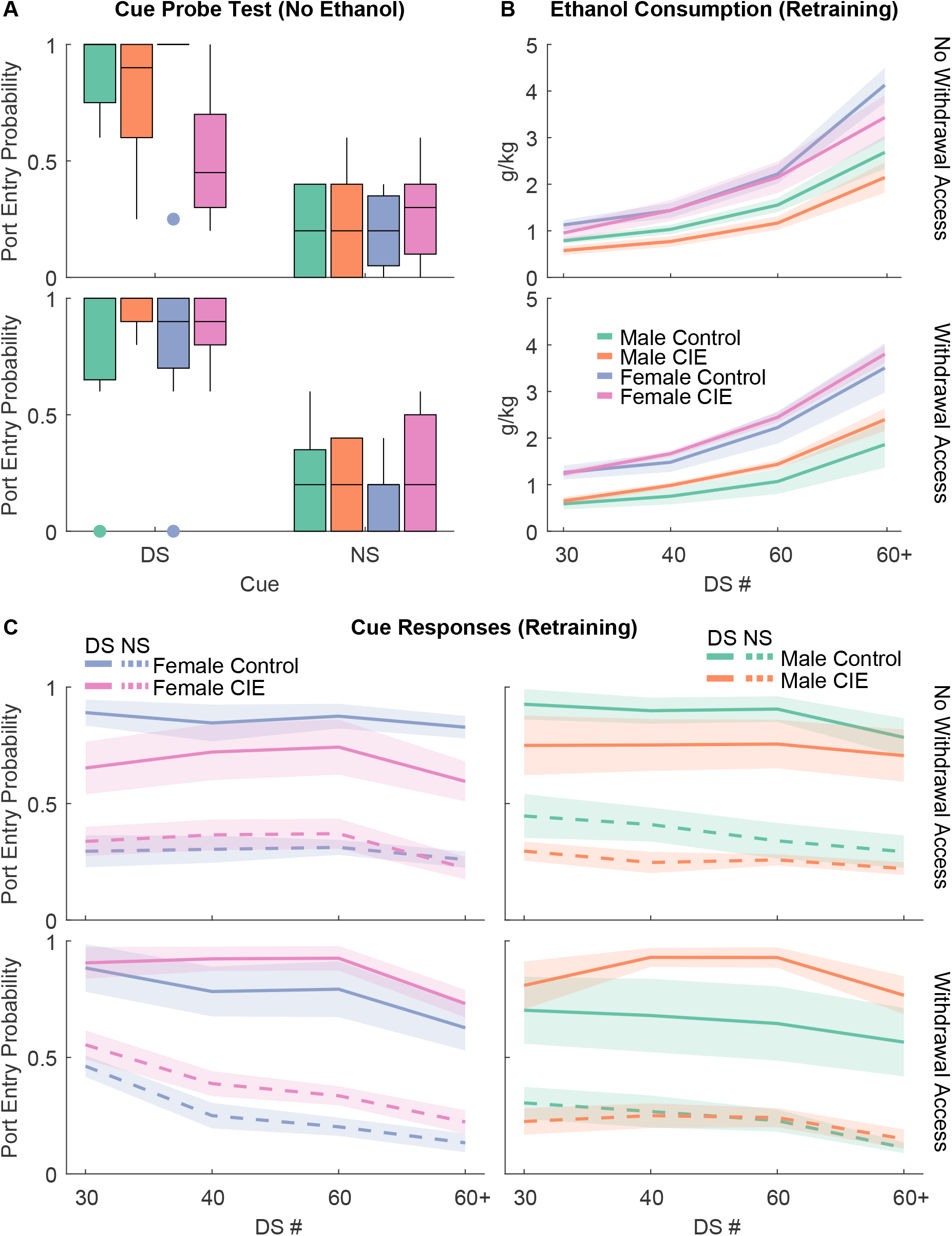
Cue responses and ethanol consumption in the DS task after vapor exposure. A) Port entry probability during the DS (left) or NS (right) in the cue probe test (no ethanol delivered) after being exposed to vapor (CIE) or control conditions. Rats are split into those that did not receive alcohol access during acute withdrawal in the vapor phase (top) or rats that did receive access in acute withdrawal (bottom). B) Ethanol consumption during cue “retraining” in which ethanol was re-introduced, and the number of trials (30-60) or the volume of reward (+) was increased across sessions. C) Port entry probability during the DS (solid lines) or NS (dotted lines) during the retraining sessions.

### CIE vapor inhalation reduced entries into the alcohol port during subsequent cue learning in male rats

While CIE vapor inhalation appeared to enhance responsivity to previously learned alcohol cues in rats that had access to ethanol during acute withdrawal, the effect size was small to moderate (Cohen’s d = 0.4632). Therefore, we next sought to determine whether the induction of alcohol dependence with CIE vapor inhalation would enhance learning for or responses to subsequently learned alcohol cues. Rats underwent 2 weeks of homecage intermittent access to 15% ethanol and were assigned to matched CIE versus control groups. Rats underwent 3 weeks of CIE vapor inhalation as in Experiment 1 and were trained in the DS task (Fig. 3A-C). Similar to Experiment 1, females consumed more g/kg ethanol during training (Fig. 3B; main effect of sex: F(1,739)=4.85, p = 0.028; session X sex interaction, F(1,739)=16.641, p < 0.001). We found no effect of CIE or interaction with session or sex for ethanol consumption during training (F values 0.085 to 1.69). As expected, port entry probability increased across sessions (Figure 3A; F(1,739)=75.134, p < 0.001), but this effect depended on sex (session X sex interaction, F(1,739)=18.059, p < 0.001). We found a trend towards an interaction between session, sex and CIE vapor inhalation (F(1,739)=3.29, p = 0.070), though this interaction did not reach statistical significance. Overall, CIE rats appeared to have blunted port entry probability during the DS compared to controls. This decrease was not specific to the DS. Port entry probability during the NS was blunted by CIE (main effect of CIE, F(1,400)=5.3685, p = 0.021; session X CIE interaction, F(1,400)=2.9211, p = 0.088). We found no effect of sex or interaction with CIE or session for NS port entry probability (F values 0.0071 to 1.49). Additionally, we found that total port entries during DS training differed based on CIE vapor exposure (Figure 3C; CIE, F(1,739)=8.2778, p = 0.0041), sex (F(1,739)=9.0935, p = 0.00265) and potentially their interaction (CIE X Sex, F(1,739)=2.844, p = 0.092). Male CIE animals had blunted port entries overall relative to controls (main effect of CIE, F(1,391)=8.9437, p = 0.00296; session X CIE interaction, F(1,391)=10.29, p = 0.0014), but female CIE animals did not (main effect of CIE, F(1,348)=0.13, p = 0.71; session X CIE interaction, F(1,348)=0.022, p = 0.88). Overall, CIE exposure did not enhance cue learning or responsivity during training and appeared to generally reduce entries into the alcohol port, especially in male rats.

**Figure 3.**
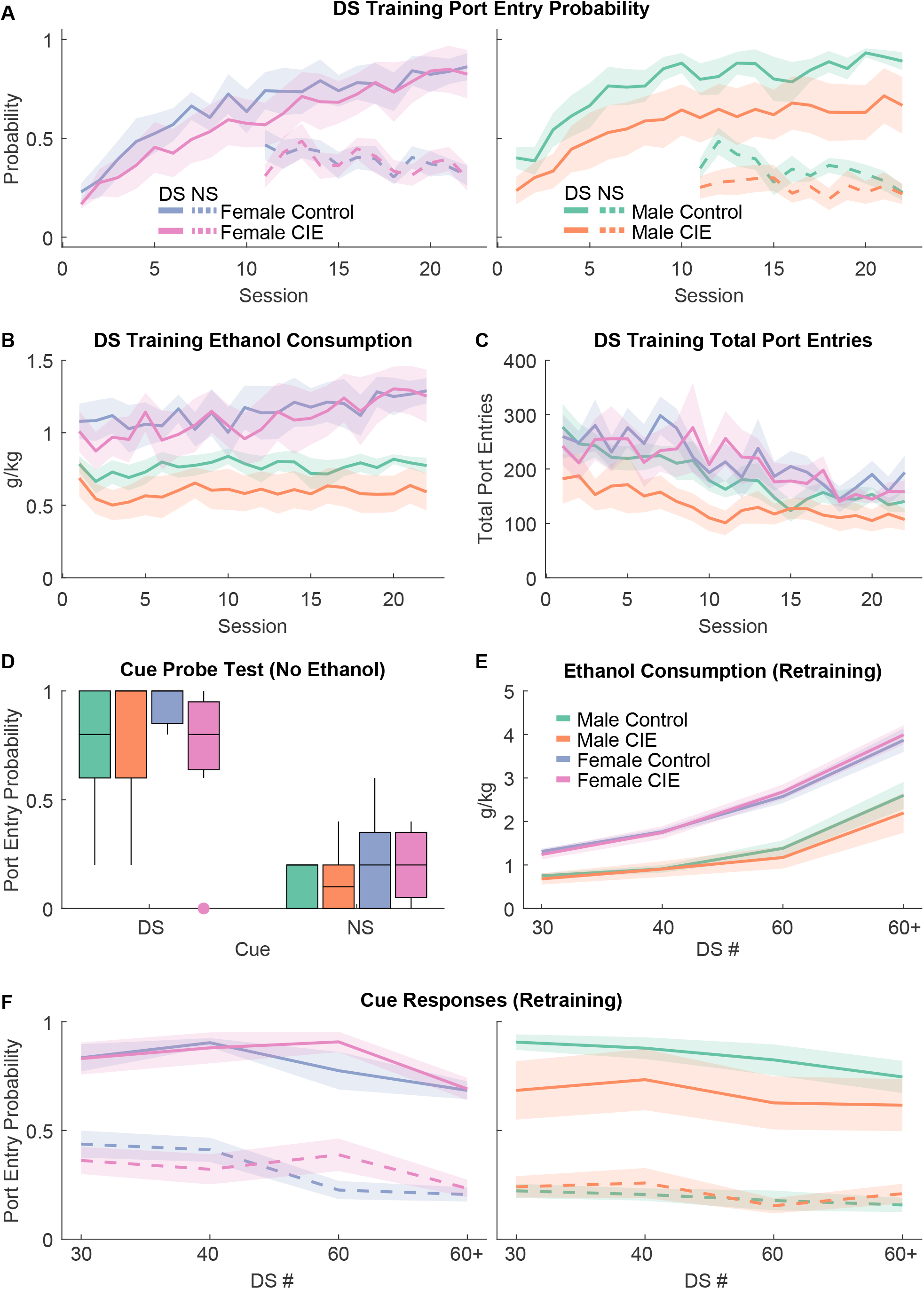
DS task training after vapor exposure. A) Port entry probability during the DS (solid lines) or NS (dotted lines) during DS task training after exposure to ethanol vapor (CIE) or control conditions. B) Ethanol consumption (g/kg) during DS task training after CIE or control conditions. C) Total port entries during DS task training after CIE or control conditions. D) Port entry probability during the DS (left) or NS (right) in the cue probe test (no ethanol delivered) that occurred after DS task training. E) Ethanol consumption during cue “retraining” in which ethanol was re-introduced, and the number of trials (30-60) or the volume of reward (+) was increased across sessions, in rats that received vapor exposure (CIE) or control conditions prior to task training. F) Port entry probability during the DS (solid lines) or NS (dotted lines) during the retraining sessions.

After another set of drinking tests, rats underwent a cue probe test and retraining testing as conducted in Experiment 1 (Fig. 3D-F). During the cue probe test (Fig. 3D) we found no effect of CIE or sex, or any interaction for DS response probability (Fs 0.23 to 0.69), DS response latency (Fs 0.09 to 0.32), or NS response probability (Fs 0.61 to 2.38). We observed a trend towards an effect of sex on total port entries (F(1,25)=3.43, p = 0.078), but no effect of CIE or interaction (Fs 0.28 to 0.1). During retraining we found a significant interaction between sex, CIE vapor inhalation, and test number for ethanol consumption (F(1,96)=5.47, p = 0.021), and a significant interaction of sex and CIE for DS response probability (F(1,96)=10.446, p = 0.0017), but no effect of CIE on NS response probability (F(1,96)=2.32, p = 0.13), or interactions of CIE with other factors for this behavioral measure (Fs 0.33 to 1.95). In male rats we found that CIE significantly reduced DS response probability across the retraining sessions (main effect of CIE, F(1,44)=12.149, p = 0.0011), but did not significantly alter NS response probability (F(1,44)=2.92, p = 0.095) or total port entries (F(1,44)=2.12, p = 0.15). In females we did not find a significant effect of CIE on DS response probability (main effect of CIE, F(1,52)=0.24, p = 0.63; CIE x test interaction, F(1,52)=0.015, p = 0.90). Overall, CIE vapor inhalation prior to DS task training appeared to reduce alcohol port entry behavior during training, as well as DS-evoked port entry behavior during retraining with ethanol, in male but not female rats. Prior CIE vapor inhalation had no effect on cue-evoked behavior in the absence of alcohol delivery.

### CIE vapor inhalation produced high blood ethanol concentrations in males and females, but only increased alcohol consumption in male rats

During the 3-week CIE vapor inhalation period, blood ethanol concentrations (BECs) were assessed immediately after the first vapor exposure session of the week (Fig. 4A). As expected, we observed high BECs (median value between 200-300 mg/dl) that did not differ between male and females (F(1,93)=0.587, p = 0.44), though we did see a trend towards an effect of week (F(1,93)=9.8357, p = 0.053). During the CIE period, we also assessed ethanol drinking behavior 2-times a week in 2-hr sessions occurring 6-8 hours after CIE rats were removed from vapor chambers. Overall, we found that CIE vapor inhalation drove increased consumption across test sessions (CIE x test interaction, F(1,347)=7.85, p = 0.0054). Because we observed an interaction of sex with test session and the timing of vapor exposure (Experiment 1 versus Experiment 2; F(1,347)=4.65, p=0.032) and a trend towards an interaction of these factors with CIE (F(1,347)=3.18, p = 0.075) we split males and females for follow-up analyses. Overall, CIE male rats increased alcohol consumption relative to controls (Fig. 4B, bottom; test X CIE interaction, F(1,166)=6.87, p = 0.0096), but female CIE rats did not (Fig. 4B, top; main effect of CIE and interactions with other factors, Fs <0.58), consistent with some prior reports (Morales et al., 2015).

**Figure 4.**
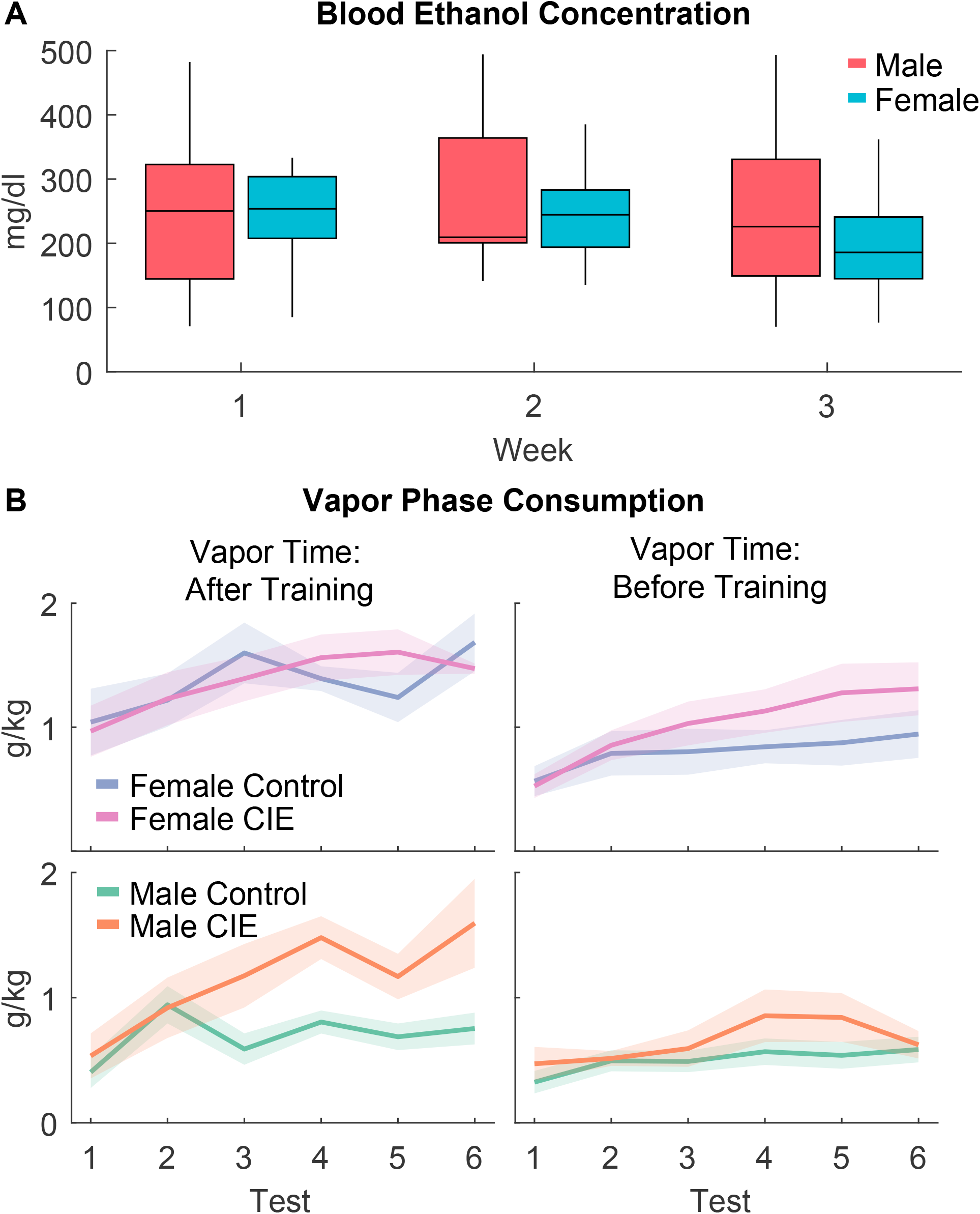
Blood ethanol levels and ethanol consumption during the vapor exposure phase. A) Blood ethanol concentrations (BECs) measured after the first vapor exposure of the week (Tuesday mornings) across three weeks of vapor exposure. B) Ethanol consumption (g/kg) during the twice weekly ethanol consumption tests 6-8 hours after CIE rats were removed from the vapor chambers. Rats on the left received vapor exposure (CIE) or control conditions after DS task training. Rats on the right received CIE or control conditions prior to DS task training.

### CIE vapor inhalation increased somatic withdrawal signs, but not thigmotaxis

To assess anxiety-like and locomotor behavior, rats were monitored in an open field prior to CIE (Test 1), at the end of CIE during acute withdrawal (Test 2), and 2 weeks after the completion of CIE (Test 3; Fig. 5A-B). When we examined mean distance from center (Fig. 5A), we found a significant interaction between CIE, sex and CIE time (before or after DS task training; F(1,222)=4.53, p = 0.034) and a trend towards an interaction of these factors with test (1,222)=3.1377, p = 0.0774). When we just examined rats that had received CIE after training in the DS task (‘After Training’; Experiment 1) we found a significant interaction between CIE and test (F(1,151)=4.96, p = 0.027), though the effect appeared to be in the opposite of the expected direction: CIE rats had decreased distance from center relative to controls. In rats that received CIE or control conditions prior to task training (‘Before Training’; Experiment 2) we found a significant effects of CIE group (F(1,71)=4.2, p = 0.044), sex (F(1,71)=7.0861, p = 0.009) and an interaction of CIE and sex (F(1,71)=3.96, p = 0.050), which appeared to be driven by pre-existing group differences, as we found no interaction of test with CIE and sex (F(1,71)=1.90, p = 0.17). Locomotor activity (distance traveled; Fig. 5B) decreased across the test sessions (main effect of test, F(1,220)=22.217, p < .001). We observed no effect of CIE on distance traveled, or interactions of CIE with vapor time, sex or test number (Fs 0.05 to 1.74). Overall, we did not find any evidence of increased anxiety-like behavior in the open field in CIE rats, and, if anything, CIE rats spent their time closer to the center. In contrast, we observed increased signs of somatic withdrawal in rats exposed to CIE vapor inhalation in a test conducted 6-8 hours after vapor exposure during the final week of CIE (Fig. 5C; main effect of CIE, F(1,69)=5.6197, p = 0.02). We did not find a significant interaction of CIE and sex (F(1,69)=2.554, p = 0.1145) or any other main effects or interactions of other factors (Fs < 0.01 to .834).

**Figure 5.**
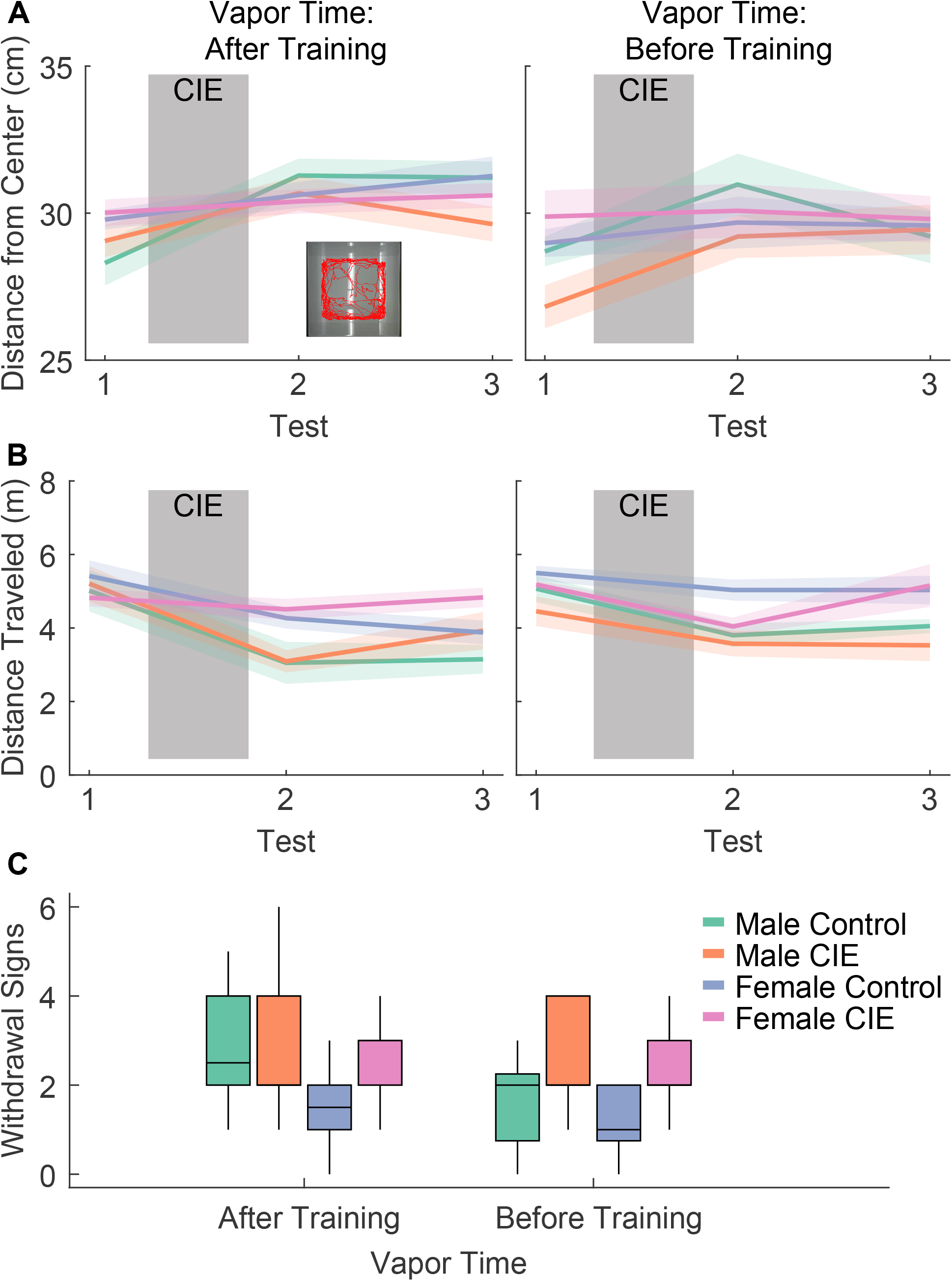
Open field behavior and somatic withdrawal symptoms. A) Distance from the center of an open field before (Test 1), the day after (Test 2) and 2 weeks after (Test 3) exposure to ethanol vapor (CIE) or control conditions. Rats on the left received vapor exposure (CIE) or control conditions after DS task training. Rats on the right received CIE or control conditions prior to DS task training. B) Distance travel in the open field during the three tests. C) The number of somatic withdrawal signs observed 6-8 hours after vapor exposure during the final week of the vapor phase.

### Sex-dependent effects of CIE vapor inhalation on aversion-resistant drinking

To assess “aversion-resistant” drinking before (Experiment 1 and 2) and after (Experiment 2) CIE vapor inhalation we measured consumption of 15% alcohol with and without quinine adulteration (Fig 6). Prior to CIE we found a main effect of quinine (Fig 6A-B; F(1,277)=5.9669, p = 0.015), which reduced ethanol consumption as expected. Rats did not differ based on their CIE group assignments (F(1,277)=2.0791, p = 0.15) and we found no interaction of quinine concentration and sex (F(1,277)=0.373, p = 0.54). In experiment 2 we re-tested rats after CIE vapor inhalation (Fig. 6C-D, left) and found a significant interaction between quinine concentration, CIE and sex (F(1,151)=5.5964, p = 0.0193). When we split males and females for separate analyses we found a significant effect of quinine concentration (F(1,110)=31.63, p < 0.001) and interaction of concentration and CIE (F(1,110)= 4.32, p = 0.04) in males, but only an effect of quinine concentration in females (main effect of quinine; F(1,97)=10.91, p = 0.001; quinine X CIE, F(1,97)=1.66, p = 0.20), suggesting that CIE vapor inhalation produced aversion-resistant drinking in males but not females. When rats were retested after training in the DS task, we found significant effects of quinine concentration (F(1,96)=9.60, p = 0.0026), CIE vapor inhalation (F(1,96)=4.09, p = 0.46) and sex (F(1,96)=13.66, p < 0.001) on g/kg ethanol consumption, but no interactions between any of these factors (Fs 0.10 to 2.58), indicating that the male-specific effects of CIE on aversion resistance did not persist after task training.

**Figure 6.**
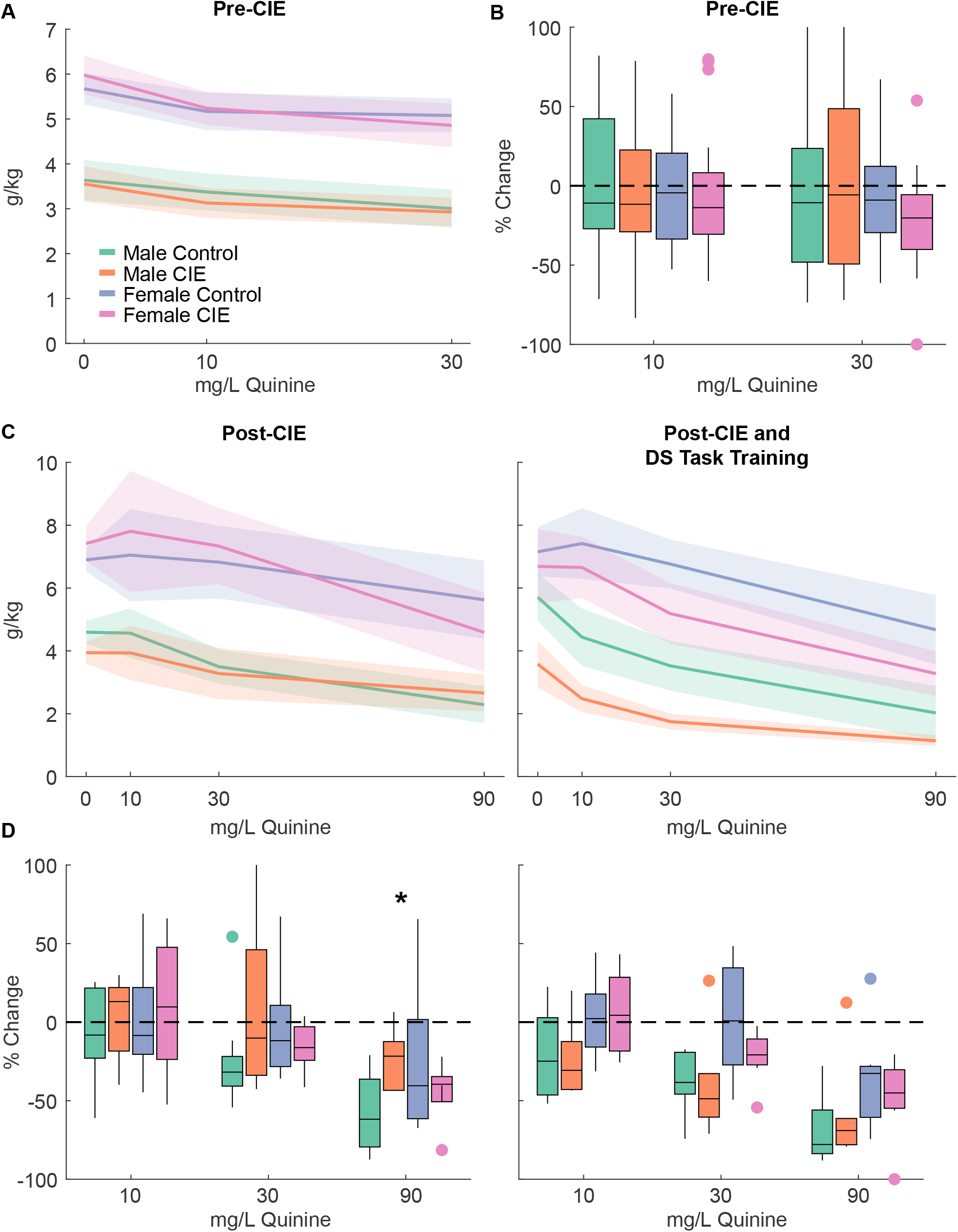
Quinine-adulterated alcohol consumption. A) Mean ethanol consumption (g/kg) with different concentration of quinine prior to exposure to ethanol vapor (CIE) or control conditions. B) Boxplots of the % change in consumption from ethanol alone at 10 and 30 mg/L quinine. C) Mean ethanol consumption (g/kg) with different concentration of quinine after exposure to ethanol vapor (CIE) or control conditions (left) and after subsequent DS task training (right). D) Boxplots of the % change in consumption from ethanol alone at 10, 30 and 90 mg/L quinine after exposure to ethanol vapor (CIE) or control conditions (left) and after subsequent DS task training (right). * = p < 0.05 interaction between CIE vapor inhalation and sex.

## Discussion

Here, we demonstrate that CIE vapor inhalation can enhance cue-elicited alcohol-seeking behavior, but only under specific circumstances. When rats were exposed to CIE after cue learning and were allowed to repeatedly access oral alcohol while in acute withdrawal during the vapor phase, we later saw an increase in the probability of alcohol port entries during the cue, under extinction and retraining conditions. In contrast, rats that did not receive access to alcohol during acute withdrawal did not show an increase in their responses to alcohol cues. Additionally, rats that received CIE vapor inhalation prior to cue learning did not show enhanced responses to cues, and male rats in particular engaged in less alcohol-seeking behavior during the task. While CIE increased responses to previously learned alcohol cues in both male and female rats that received acute withdrawal access, we observed sex-specific effects on some other measures. Specifically, CIE vapor inhalation increased alcohol consumption and aversion-resistant drinking selectively in male rats.

### Sex differences in ethanol consumption and the effects of CIE vapor inhalation

In the current studies we found that females consumed more g/kg ethanol during homecage intermittent access and DS task training, and that CIE vapor inhalation increased ethanol consumption and “aversion-resistant” drinking in male, but not female rats. Elevated ethanol consumption in female rats is consistent with a rich prior literature documenting sex differences in alcohol consumption in rodents (Becker and Koob, 2016). Female rats, including Long Evans, have been shown to consume more g/kg ethanol under both continuous (Juárez and De Tomasi, 1999; Lancaster and Spiegel, 1992; Li and Lumeng, 1984; Lorrai et al., 2019) and intermittent access conditions (Aguirre et al., 2020; Priddy et al., 2017), as well as during self-administration (Blanchard et al., 1993; Randall et al., 2017). Whether these differences in voluntary consumption result in different blood ethanol concentrations in female and male rats remains unclear. Some studies have reported that males have higher blood ethanol concentrations after receiving identical doses of injected ethanol (Mankes et al., 1991; Morales et al., 2015; Rachamin et al., 1980), but others have reported similar blood ethanol levels (Randall et al., 2017). Blood ethanol levels have been reported to be similar at the end of a 4 hour drinking session (Marco et al., 2017), but these values likely do not correspond to peak levels during these sessions. Whether female rats are consuming more g/kg to reach similar or greater blood ethanol concentrations in comparison to male rats remains an important open question.

Our finding that CIE vapor inhalation increases alcohol consumption in male but not female rats is also consistent with some prior work (Morales et al., 2015; but see Priddy et al., 2017). This may be due in part to the heightened consumption in control females; CIE vapor inhalation drives consumption in male rats that is similar to consumption levels in control females. We also found that CIE vapor inhalation selectively impacted aversion-resistant drinking in male, but not female rats. We found no impact of sex independently of CIE vapor inhalation, consistent with some prior reports (DeBaker et al., 2020; Randall et al., 2017; Sneddon et al., 2018), but not others (Fulenwider et al., 2019; Sneddon et al., 2020). Prior reports examining the impact of CIE vapor inhalation on aversion-resistant drinking have not, to our knowledge, included female subjects (Vendruscolo et al., 2012), though differences in other measures of compulsive alcohol seeking have been reported (Xie et al., 2019). Despite these sex differences in baseline intake and the effects of CIE vapor inhalation on intake and aversion-resistant drinking, we saw similar effects of CIE vapor inhalation on behavioral responses to previously-learned ethanol cues in males and females. Yet, given the other sex differences we found, and the long history of male-only CIE vapor inhalation studies, it is important to consider whether prior effects reported for CIE vapor inhalation are generalizable to female rodents. Additionally, even when similar behavioral effects are reported, these effects may be driven by different psychological and neurobiological mechanisms.

### Psychological mechanisms underlying CIE vapor inhalation effects on cue responsivity

The importance of access to alcohol during acute withdrawal for the cue effects reported here may shed some light on the potential psychological mechanisms underlying these CIE vapor inhalation effects. Increased alcohol consumption and seeking after CIE vapor inhalation has generally been attributed to activation of stress systems and/or negative reinforcement mechanisms (de Guglielmo et al., 2016; Gilpin and Koob, 2010; Koob et al., 2014; Somkuwar et al., 2017; Tunstall et al., 2017; Vendruscolo et al., 2012; Walker, 2012). While these explanations are frequently discussed in combination, they are not synonymous. Stress can also potentiate cue salience and responsivity without the opportunity for new learning via negative reinforcement (Glynn et al., 2018; Sinha and Li, 2007). Our results suggest that new learning about alcohol is necessary for potentiated cue responding following CIE vapor inhalation, but we cannot rule out that a shorter delay between CIE vapor inhalation and cue testing might yield different results. While our observed changes in cue-elicited behavior were gated by access to alcohol during acute withdrawal, they were not dependent on re-experiencing the cue-alcohol association, either during acute or protracted withdrawal. Instead, the patterns we observed during retraining and extended cue testing were already present during our initial cue probe test. This suggests that reappraisal of the value of the alcohol during acute withdrawal may transfer to the cue itself without additional cue-alcohol pairings.

### Learning and responsivity to non-alcohol cues is impacted by CIE vapor inhalation

Under some conditions, rather than observing increases in cue responsivity, we actually saw decreases in cue reactivity and/or alcohol seeking. These findings are less surprising in the context of prior work on behavioral responses to non-alcohol cues. Deficits in learning, discrimination or reactivity to non-alcohol cues have been reported after CIE vapor inhalation (Barker et al., 2016), especially after more prolonged CIE procedures in both rats and mice (DePoy et al., 2015; Natividad et al., 2018). Yet, CIE vapor inhalation has also been reported to enhance subsequent responding to non-alcohol cues by mice, including in tests of Pavlovian-to-instrumental transfer (Shields and Gremel, 2021), as well as pairwise visual discrimination and reversal learning (DePoy et al., 2013). At this point it is difficult to reconcile these disparate findings, but it is likely that both the duration and time since CIE vapor inhalation impact the direction of the effects, with longer and/or more recent exposure being more likely to produce deficits in cue-guided behavior and memory.

## Conclusions

Here we found that CIE vapor inhalation potentiated responding to previously learned alcohol cues when rats had the opportunity to consume alcohol during acute withdrawal, but that it failed to enhance cue responding either in the absence of this access or when cue-alcohol training occurred after CIE. This suggests that high levels of alcohol exposure alone are not enough to produce the types of behavioral changes that result in heightened risk of relapse and craving even after long periods of abstinence. Instead, the learning that occurs during voluntary alcohol consumption and self-administration is a critical factor in the degree to which alcohol cues can control behavior. Additionally, sex differences in the ability of CIE to alter drinking, but not cue-evoked responding, indicate that altered consumption is not a prerequisite for altered alcohol cue reactivity. Together these results highlight the importance of selecting appropriate alcohol seeking and exposure models, tested in both male and female subjects, to better understand the neural and psychological mechanisms underlying alcohol addiction.

## Acknowledgements

This work was supported in part by National Institutes of Health grants R00AA025384 and R01AA028770 to JMR.

## Notes

### Competing Interest Statement

The authors have declared no competing interest.

